# Detection of Merkel cell polyomavirus using whole exome sequencing data

**DOI:** 10.1101/2020.04.27.063214

**Authors:** Sandra Garcia-Mulero, Ferran Moratalla-Navarro, Soraya Curiel-Olmo, Victor Moreno, José Pedro Vaqué, Rebeca Sanz-Pamplona, Josep María Piulats

## Abstract

Merkel cell carcinoma (MCC) is a highly malignant neuroendocrine tumor of the skin in which Merkel cell polyomavirus (MCV) DNA virus insertion can be detected in 75–89% of cases. Etiologic and phenotypic differences exist between MCC tumors with and without the inserted virus, thus it is important to distinguish between MCV+ MCC and MCV-MCC cases. Currently, MCV insertions in MCC genomes are detected using laboratory techniques. Here we report a freely available bioinformatics methodology to identify MCV+ MCC tumors using whole exome sequencing (WES) data. WES data could be also used to infer the virus insertion site into the tumor genome. Our method has been validated in a set of MCC samples previously characterized in the laboratory as MCV+ or MCV-, achieving 100% sensitivity and 62,5% specificity. Thus, with enough depth of sequencing, it is possible to use WES to the presence of MCV insertions in cancer samples.

## Introduction

Merkel cell carcinoma (MCC) is highly malignant neuroendocrine tumor of the skin currently thought to arise from mechanoreceptor Merker cells [1]. MCC typically affects immunosuppressed individuals thus suggesting an infectious origin. Indeed, in 2008, a 5.4 kbp polyomavirus DNA was found to be integrated in some MCC genomes in a clonal pattern [2]. Although MCC is rare, its incidence has tripled over the past two decades in the Unites States. This could be due for both the increasing prevalence of risk factors such as ultraviolet (UV) exposure or systemic immune suppression [3] or because of the improvement of diagnostic methods. Merkel cell polyomavirus (MCV) DNA virus can be detected in 75–89% of MCC. It is noteworthy that etiologic and phenotypic differences exist between MCC tumors with and without the inserted virus [4]. In this regard, recent work by independent laboratories has shown important genetic differences between MCV+ and MCV-MCC tumors, the latter harboring higher mutational burdens with ultraviolet (UV) signatures [5–8]. Moreover, relevant clinical features are associated with this phenotype being the MCV-subtype the one showing a worst prognosis [9]. Also, from a therapeutic point of view, immunotherapy could play an important role in this tumor [10, 11]. Thus, it is important to distinguish between MCV+ MCC and MCV-MCC. Normally, MCV insertions in MCC genomes were detected by PCR or using an anti-MCP antibody [9]. Here we show a 2-alignment-steps methodology to identify MCV+ tumors using WES data.

## Results and discussion

### 2-alignment-steps pipeline

The following pipeline was set up using a positive control sample (WES data from a MCV+ MCC tumor analyzed with standard laboratory techniques): after quality control assessment using FastQC [12], WES raw data was pre-processed with TrimGalore [https://github.com/FelixKrueger/TrimGalore] for adapters and bad quality reads removal. Then, reads were aligned against the human reference genome with Bowtie2 [13] using strict parameters to achieve a very sensitive local alignment. Next, unmapped reads (reads that did not map to human genome) were retrieved using SamTools [14]. Finally, these non-human reads were aligned to the NCBI Merkel cell polyomavirus reference genome (EU375803.1; isolate MCC350, complete genome). This second alignment was performed with BWA aligner [15]. If reads were found matching with MCV genome, the tumor was classified as MCV+ (Figure 1). In detail description of the pipeline is found in the Materials and Methods section. The script used to make this analysis is freely available at Github repository (https://github.com/odap-ubs/merkel_virus). We had previously assessed that there was no similarities between the MCV genome and any sequence in the Homo sapiens genome. The MCV genome is about 5Kb long, and splits of 75bp (a total of 70) were submitted to the online tool BLAST [16]. No homology was found. Per sample coverage was calculated as twice the read length multiplied by total number of reads and divided by total genome length.

**Figure 1.**
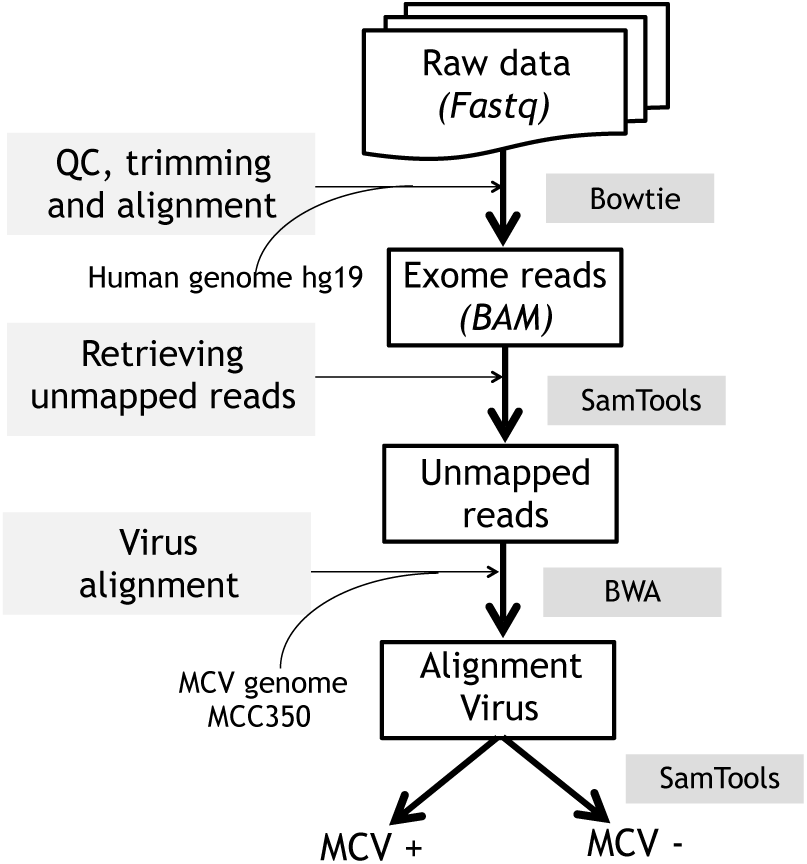
Pipeline for Merkel virus insertion detection using WES data.

### MCV insertion in MCC patients

We interrogated sequencing data for a total of 15 MCC primary tumors and their normal paired samples, previously characterized in the laboratory for MCV insertions (seven MCV+, and eight MCV-) [5]. Our method was able to detect all seven MCC positive for MCV (ranged from 2 to 44 mapped reads). Supplementary Figure 1 shows the MCV+ MCC aligned reads in the Integrative Genome Viewer (IGV) [17]. Interestingly, there was a tendency towards detecting more aligned reads in the first 2,500 bp of the MCV. As expected, none of the normal samples were found to present MCV insertions. Strikingly, three out of eight previously classified as MCV-MCC were found to have at least one read mapping into MCV genome whereas the remaining five had zero mapped reads. Sensitivity and specificity were 100% and 62.5%, respectively. However, due to the extremely low number of reads in false positive samples, better values of specificity would be obtained if a sample was catalogued as positive only when at least two MCV reads were detected. It was also interesting to note that there was a correlation between the sequencing coverage and the number of mapped reads. This may indicate that when depth of sequencing increases also increases the probability of finding an inserted viral genome (Table 1).

**Table 1.** Alignment results.

| SAMPLE | TISSUE | MCP | N° MAPPED READS | COVERAGE |
| --- | --- | --- | --- | --- |
| 48120427 | FF | + | 44 | 3.519 |
| 48120428 | FF | + | 34 | 2.704 |
| 48121209 | FF | + | 16 | 2.543 |
| 48120987 | FF | + | 15 | 2.286 |
| 48140221 | FF | + | 7 | 2.090 |
| 48090369 | FF | + | 4 | 2.467 |
| 48130247 | FFPE | + | 2 | 1.794 |
| 48130206 | FFPE | - | 2 | 1.304 |
| 48120431 | FF | - | 1 | 3.213 |
| 48141029 | FFPE | - | 1 | 1.784 |
| 48141028 | FFPE | - | 0 | 1.869 |
| 48130208 | FFPE | - | 0 | 1.364 |
| 48130207 | FFPE | - | 0 | 1.691 |
| 48121576 | FF | - | 0 | 2.520 |
| 48120426 | FF | - | 0 | 3.192 |

Since interrogated samples were a mixture of fresh frozen (FF) and formalin-fixed paraffin-embedded (FFPE) preserved tumors, we wondered if our method performed well in both types of samples. Figure 2 suggest that our pipeline is more robust when NGS was performed in FF samples. This is not surprising since sequencing performs better in FF tissues. However, only one out of seven MCV+ MCC were FFPE. Moreover, the MCV+ FFPE sample was the one with the lower sequencing depth so a coverage bias rather than a tissue preservation effect cannot be excluded. Thus, more samples need be assessed in order to settle this topic.

**Figure 2.**
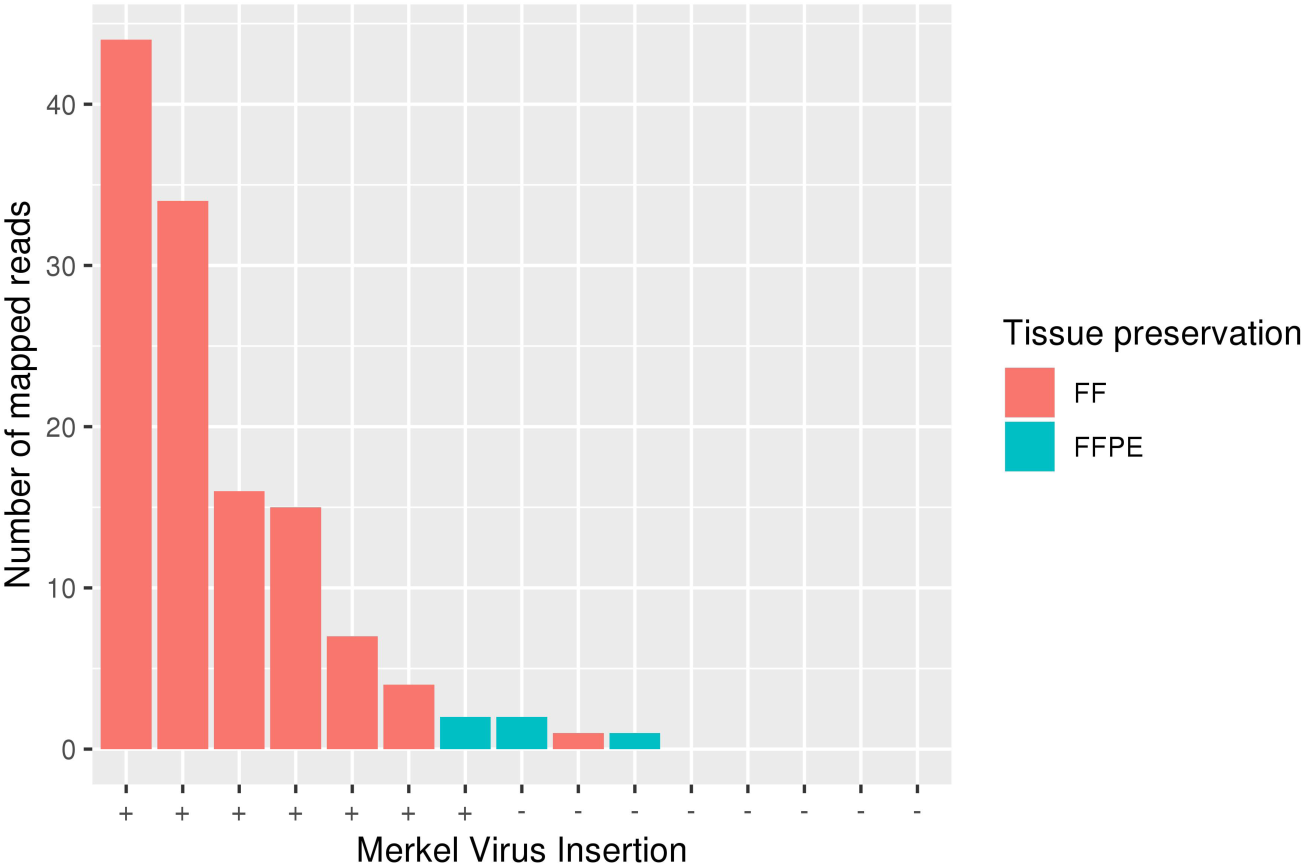
Barplot representing the number of mapped reads by sample. Fresh frozen (FF) samples were represented in orange and formalin fixed paraffin embedded (FFPE) in green. In the x axis, positive (+) and negative (-) MCC tumors were indicated.

### MCV site of insertion

Finally, paired-end WES data was used to try to infer the virus insertion site into the tumor genome. After applying this strategy in all positive samples, only one of them (48121209) showed soft-clipped reads whose mates were mapped on chr19:48,445,990. Interestingly, this region was into GRWD1 gene intronic region (Supplementary Figure 2A). This is a P53 regulator whose loss of function has been previously associated with tumorigenesis [18]. The fact that this is a highly covered region might indicate that our method failed to detect insertion positions in poor covered regions. Thus, whole-genome sequencing should be a better technique to identify not only MCV+ tumors but also the virus insertion site. Supplementary Figure 2B showed how soft-clipped reads also mapped against the MCV genome.

Apart from MCC, MCV insertions have been found in small-cell lung cancer in non-smokers [19]. However, this is a controversial result since other study has reported no MCV insertions in this cancer [20]. Moreover, MCV DNA fragments could be detected in the lower respiratory tract when a high-sensitive PCR assay was used, thus false positives could not be excluded [21]. Therefore, the 2-alignment-steps methodology was applied on non-smokers lung adenocarcinoma data sets; 25 RNA-seq samples [22] and 30 WES samples [23], but no MCV+ tumor was found.

## Conclusions

In conclusion, here we present an easy method to detect MCV insertions as a bystander product of exome sequencing. Since, human gene exons DNA are enriched in WES, MCV will be most likely detected in MCC if its genome is inserted in exome regions or when high sequencing depth is obtained and MCV reads are detected as off-target reads. We cannot rule out the possibility that samples were contaminated with viral DNA, but the lack of positive results in paired normal samples sequenced with similar coverage was reassuring. Thus, with enough depth of sequencing, it is possible to apply the pipeline described here to take advantage of WES experiments and assess the presence of MCV insertions in any dataset of interest.

## Materials and Methods

### Patients and samples

WES data from a total of 15 MCC patients and their normal paired samples were used for the analysis. Samples were either formalin fixed paraffin embedded (FFPE) or freshly frozen (FF). For clinical characteristics and genomic analysis, refer to González-Vela et al. study [5]. For validation purposes, data from the European Nucleotide Archive (ENA) public repository was downloaded; 25 RNA-seq samples from series SRP090460 [22], and 30 WES samples from series SRP022932 [23] were used.

### Identification of MCV+ tumors using WES

First, raw data was evaluated for quality control with FastQC. Next, samples were trimmed if necessary with TrimGalore (v0.4.0) and aligned against the human reference genome build hg19/ GRCh37, previously indexed, using Bowtie 2.0 (v2.2.5) ─alignment 1─. From this first alignment, unmapped reads are selected with Samtools (v1.3.1). Function Samtools view, with -f and -F options, was used to select and filter the desired reads. The online utility “Decoding SAM flags” from the Broad Institute (https://broadinstitute.github.io/picard/explain-flags.html) was used to decide the filtering criteria. The selection criteria filters: i) unmapped reads whose mate are mapped (-f 4 -F 264), ii) mapped reads whose mates is unmapped (-f 8 -F 260), iii) both unpaired reads (-f 12 -F 256). The three outputs are merged and next, those retrieved unmapped reads are aligned against MCV genome (5,381 bp length, downloaded from the NCBI ─GenBank accession number EU375803.1; Merkel cell polyomavirus isolate MCC350, complete genome─) with BWA (v0.7.15) ─alignment 2─. Finally, the number of reads aligned to virus is calculated, and an output is generated with the following information: Sample ID, MCV+/− status, and read counts in case of MCV+.

### Virus insertion site

After the first alignment against the human reference genome, unmapped and discordant reads were kept to be aligned against the MCV genome. Of these, reads mapped to the MCV genome whose mates were mapped to the human genome were used to get information of the proximal insertion site (Supplementary Figure 3). On the other hand, a visual inspection of the bam files using IGV tool was done in order to find the exact site of insertion.

## Author Contributions

RSP and JMP designed the study, conceived the experiments and wrote the article. SGM and FMN carried out the sequencing analyses. SCO and JPV performed wet lab experiments. VM provides biostatistics expertise. VM and JPV helped to draft the manuscript. All authors critically reviewed and had final approval of the article.

## Acknowledgments

We thank Josipa Bilic for critical reading of the manuscript and English language assistance. We thank Spanish Association Against Cancer (AECC) Scientific Foundation.

## Supplementary Materials

**Supplementary Figure 1.**
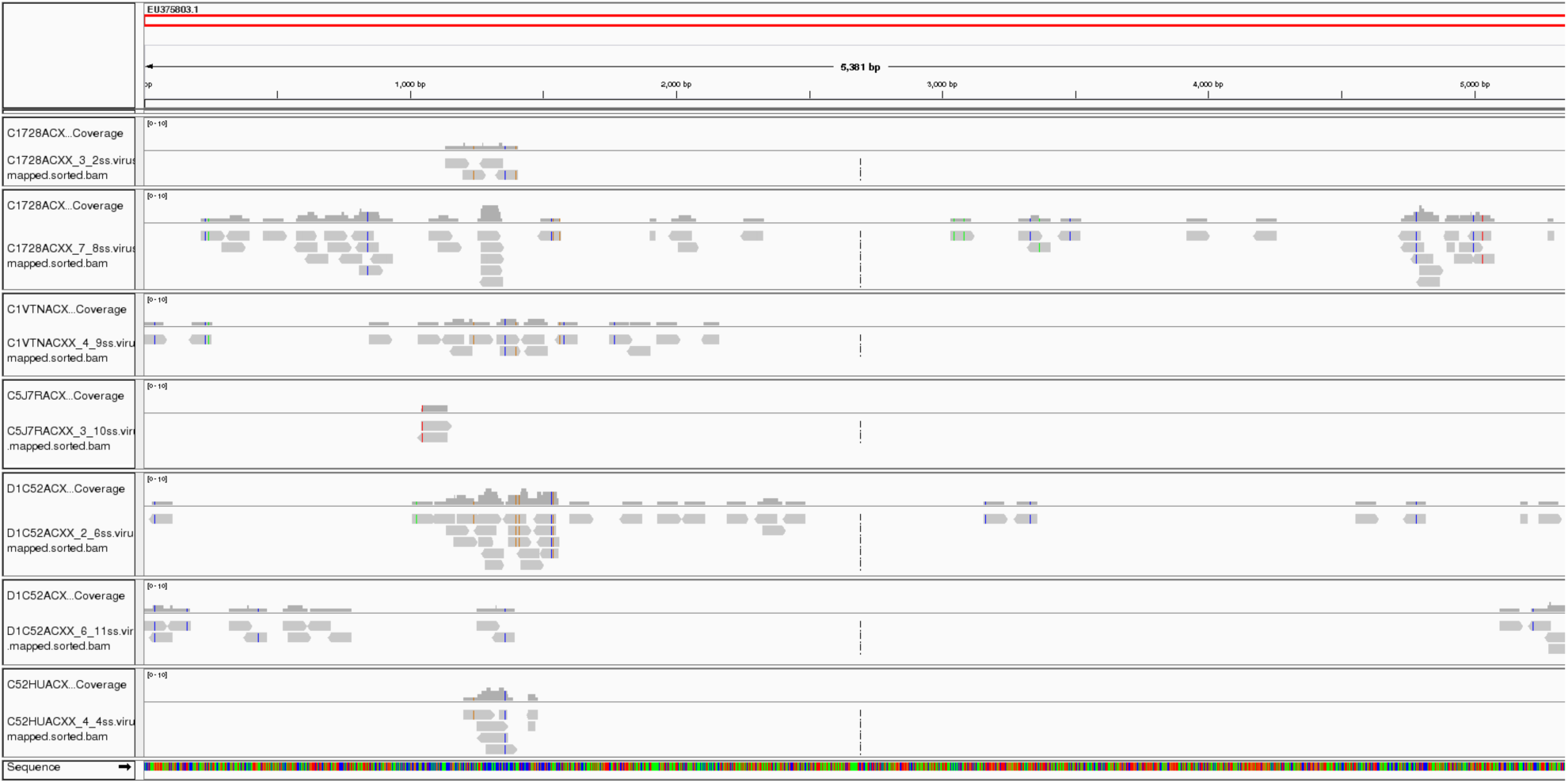
MCV+ MCC alignment IGV visualization. Upper line represented the MCV genome length. Each line corresponds to the seven MCV+ MCC tumor samples.

**Supplementary Figure 2.**
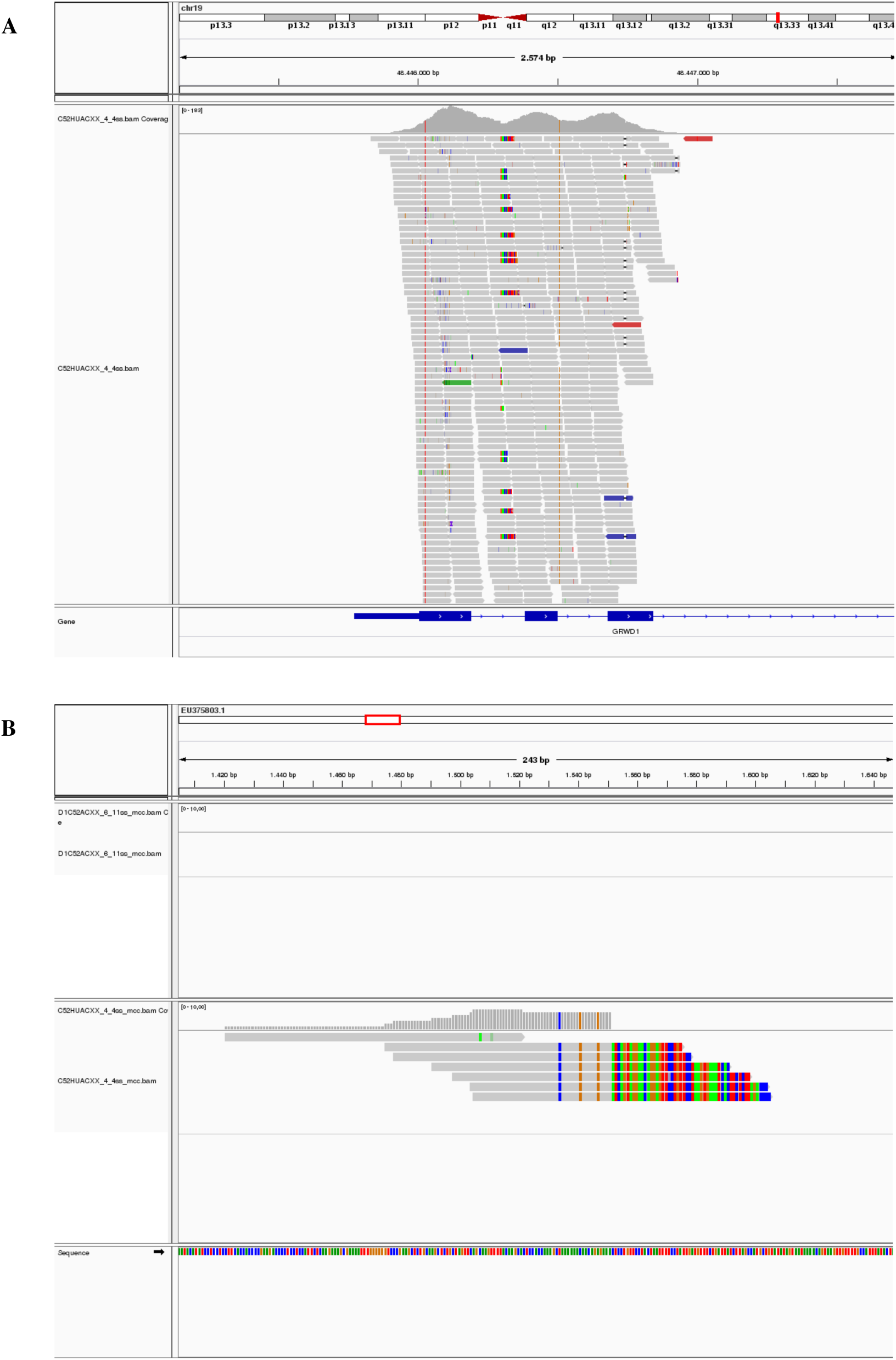
MCV insertion site in MCC sample 48121209. A. Breakpoint regions in GRWD1 gene (chromosomal coordinates chr19:48,445,990). Several soft-clipped reads in human alignment in the central part of the image, indicating the exact breakpoint in this sample. **B. Breakpoint regions in MCV.** The rainbowed part of reads (soft-clipping representation) indicates the site of the reads unable to map into MCV genome.

**Supplementary Figure 3.**
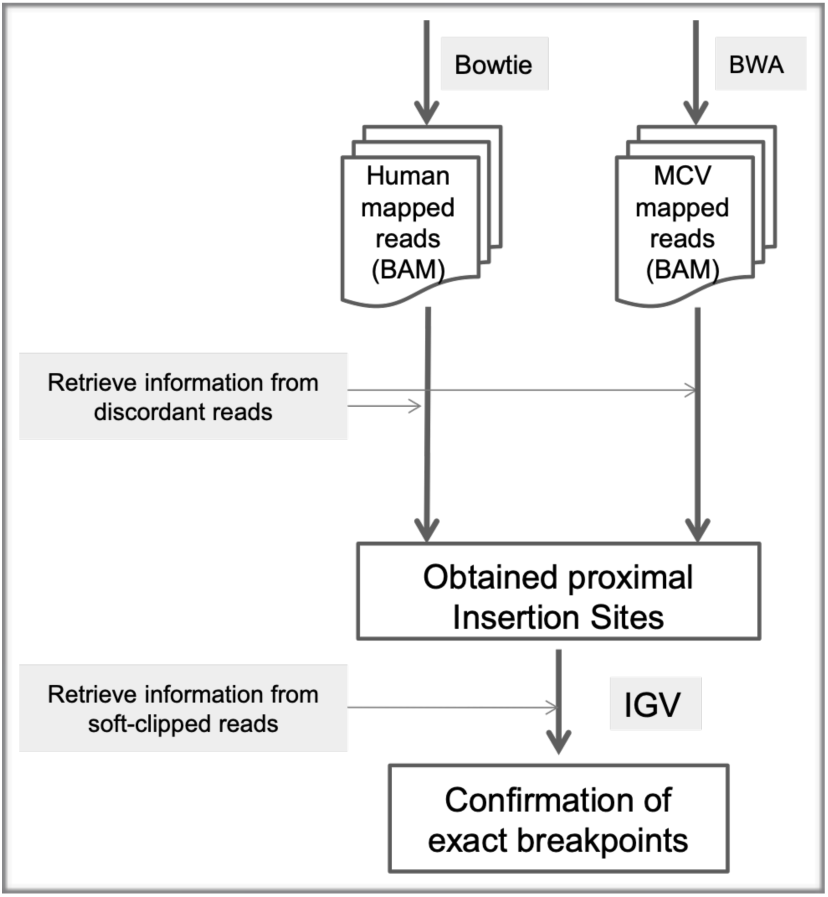
MCV insertion site into human genome pipeline. Bam files aligned into human and into MCV genome virus previously obtained were used. Discordant mapped reads of MCV alignment whose mates were mapped into human were used to narrow the possible insertion site to a few hundred of base pairs. Next, IGV was used to inspect these regions in search for soft-clipped reads both in human and virus alignment that confirmed the exact insertion site.

